# The R package *Rsubread* is easier, faster, cheaper and better for alignment and quantification of RNA sequencing reads

**DOI:** 10.1101/377762

**Authors:** Yang Liao, Gordon K. Smyth, Wei Shi

## Abstract

The first steps in the analysis of RNA sequencing (RNA-seq) data are usually to map the reads to a reference genome and then to count reads by gene, by exon or by exon-exon junction. These two steps are at once the most common and also typically the most expensive computational steps in an RNA-seq analysis. These steps are typically undertaken using Unix command-line or Python software tools, even when downstream analysis is to be undertaken using R.

We present Rsubread, a Bioconductor software package that provides high-performance alignment and counting functions for RNA-seq reads. Rsubread provides the ease-of-use of the R programming environment, creating a matrix of read counts directly as an R object ready for downstream analysis. It has no software dependencies other than R itself. Using SEQC data and simulations, we compare Rsubread to the popular non-R tools TopHat2, STAR and HTSeq. We also compare to counting functions provided in the Bioconductor infrastructure packages. We show that Rsubread is faster, uses less memory and produces read count summaries that more accurately correlate with true values. The results show that users can adopt the R environment for alignment and quantification without suffering any loss of performance.

## 1 Introduction

RNA sequencing (RNA-seq) is currently the method of choice for performing genome-wide expression profiling. One of the most popular strategies for measuring expression levels is to align RNA-seq reads to a reference genome and then to count the number of aligned reads that overlap each annotated gene [1, 2, 3]. Alternatively, reads might be counted by exon or by exon-exon junction [4]. Read mapping and read counting thus constitute a common workflow that summarizes RNA-seq reads into a count matrix, which can be used for downstream analyses such as differential expression. These two steps often represent the most computationally expensive part of an RNA-seq analysis.

R is one of the world’s most popular programming languages [5]. The TIOBE Programming Community index places it 14th overall at the time of writing and first amongst languages designed specifically for statistical analysis (https://www.tiobe.com/tiobe-index). Building on R, Bioconductor is arguably the world’s most prominent software development project in statistical bioinformatics [6]. Bioconductor contains many highly cited packages for the analysis of RNA-seq read counts, including limma [7, 8], edgeR [9] and DESeq2 [10] for differential expression analyses and DEXSeq [4] for analysis of differential splicing. Key attractions of Bioconductor include the ease-of-use of the R programming environment, the well organized package management system, the wealth of statistical and annotation resources, the interoperability of different packages and the ability to document reproducible analysis pipelines. All the RNA-seq data analysis packages rely, however, on read alignment and summarization, which typically have to be performed outside of R. This complicates the analysis pipeline, introducing additional software dependencies and creating substantial obstacles for non-expert uses.

The last decade has seen rapid development of splice-aware read alignment software. TopHat was the first successful and popular RNA-seq aligner [11]. Later aligners such as STAR [12], Subread, Subjunc [13] and HISAT [14] were dramaticaly faster while improving also on accuracy. RNA-seq read counting algorithms have developed at almost the same pace, including BEDTools [15], featureCounts [1], htseq-count [3] and Rcount [16]. htseq-count is a Python script while the other counting or mapping tools are stand-alone programs written in C or C++.

QuasR is a Bioconductor package that attempts to fill the gap, providing RNA-seq read alignment and read counting in the form of R functions [17]. QuasR is however an interface to C programs from 2010 or earlier, specifically to Bowtie version 1.1.1 [18], SpliceMap 3.3.5.2 [19] and SeqAn 1.1 [20]. These older tools do not reflect the considerable improvements in algorithms achieved during the last 8 years.

This article presents Rsubread, a Bioconductor package that implements current high-performance RNA-seq read alignment and read counting algorithms in the form of R functions. Rsubread incorporates the C programs Subread, Subjunc and featureCounts, together with other functionality. It is continuously maintained so as to track the latest versions of the C programs. Rsubread allows RNA-seq data analyses, from raw sequence reads to scientific results, to be conducted entirely in R [21, 22]. Except for read alignment itself, all Rsubread functions produce standard R data objects, allowing seamless integration with downstream analysis packages.

Many of the most popular RNA-seq algorithms are under continuous development. We take the opportunity in this article to compare Rsubread with the current versions of other tools. Previous studies have evaluated the performance of read aligners or quantifiers separately[13, 23, 1, 24]. In this article, we assess the performance of complete alignment+count pipelines and evaluate their ability to correctly represent the expression levels of genes and exons or to detect exon-exon junctions. Unlike previous studies, we consider possible interactions or incompatibilities between aligners and quantifiers.

The purpose of the article therefore is three-fold. First, we describe the functionality of the Rsubread package. Second, we demonstrate that the Rsubread is more than competitive against alternative tools, whether available in R or not, in terms of speed, memory footprint and accuracy. Third, we conduct comparisons treating read alignment and summarization together, accounting for possible interactions between aligners and read counting tools.

## 2 Materials and Methods

### 2.1 Software tools

This study compares Rsubread 1.30.5 with aligners STAR 2.6.0c and TopHat 2.1.1 and with quantifiers HTSeq 0.10.0, IRanges 2.14.10, GenomicRanges 1.32.3 and DEXSeq 1.26.0. Rsubread, IRanges, GenomicRanges and DEXSeq are R packages available from http://www.bioconductor.org. STAR and TopHat2 are Unix command-line tools written in C++ available from https://github.com/alexdobin/STAR and https://ccb.jhu.edu/software/tophat respectively. HTSeq is a Python library available from https://pypi.org/project/HTSeq.

To make a fair comparison across different workflows, aligners and quantifiers were run with similar settings as far as possible. All aligners were instructed to output no more than one alignment per read. STAR was run with the 2-pass mode. Rsubread and STAR were set to output name-sorted reads for paired end data, but TopHat2 only supports location-sorted reads. All read mapping tools were run with 10 threads. *featureCounts* is the only read counting tool that supports multi-threaded running and was run with 4 threads in the evaluation.

All timings and comparisons reported in this article were undertaken on a CentOS 6 Linux server with 24 Intel Xeon 2.60 GHz CPU cores and 512 GB of memory.

### 2.2 SEQC Data

As an example of real RNA-seq data with known expression profiles, data generated by the SEQC Project [2] was used. Two particular FASTQ files were used, one generated from sequencing of Human Brain Reference RNA (HBRR) and one from Universal Human Reference RNA (UHRR). Each file contains about 15 million 100bp read-pairs and was generated from an Illumina HiSeq sequencer.

The SEQC Project includes expression values measured by TaqMan RT-PCR for slightly over 1000 genes for both HBRR and UHRR. 958 of these TaqMan validated genes were found to have matched symbols with genes in the RNA-seq data. The TaqMan RT-PCR expression values are available from the seqc Bioconductor package.

### 2.3 Simulations

Paired-end FASTQ files were generated using the same GRCh38/hg38 genome and gene annotation as for the SEQC data. Germline variants including SNPs and short indels were introduced to the reference genome at the rates of 0.0009 and 0.0001 respectively, before sequence reads were extracted from the genome. Base substitution errors in sequencing were simulated according to Phred scores at the corresponding positions in randomly selected RNA-seq reads from an actual RNA-seq library (GEO accession GSM1819901), thus error profile of simulated reads is similar to that of real RNA-seq reads.

FPKM values were generated from an exponential distribution and randomly assigned to genes. The FPKM values were then mapped back to genewise read counts according to known gene lengths in order to achieve a library size of 15 million read pairs. Fragment lengths were randomly generated according to a normal distribution with mean 200 and variance 30. Fragment lengths below 110 or above 300 were reset to 110 or 300 respectively. Given the fragment length, a pair of 100bp sequences was randomly selected from a gene assuming that all exons for that gene are equally expressed and sequentially spliced.

### 2.4 Annotation

All evaluations and simulations used Rsubread’s built-in RefSeq gene annotation for human genome GRCh38/hg38 (build 38.2). This is identical to NCBI annotation except that overlapping exons from the same gene are merged to produce a non-overlapping set of exons for each gene. This simplification reduces ambiguity and somewhat improves the performance of all the read quantification tools. This annotation contains 28,395 genes and 261,752 exons.

### 2.5 Access to data and code

All the data and code used in this study can be freely accessed from http://bioinf.wehi.edu.au/Rsubread/. Commands for running each workflow and for producing the comparison results are included in the code.

## 3 Results

### 3.1 The Rsubread workflow

The Rsubread pipeline for read mapping and quantification consists of five R functions (Table 1). *buildIndex* builds an index of the reference genome. This is a once-off operation for each reference genome, as the same index file can be used for multiple projects. Either *align* or *subjunc* is used to align sequence reads to the reference genome and *featureCounts* produces a matrix of counts. *propmapped* is optional and produces a table of mapping statistics. In terms of output, *buildIndex*, *align* and *subjunc* write files to disk whereas *featureCounts* and *propmapped* produce R objects.

**Table 1:** The main Rsubread functions for read alignment and quantification.

| Function | Description |
| --- | --- |
| *buildIndex* | Create hash table of target genome |
| *align* | Basic alignment with soft-clipping, for gene-level analyses |
| *subjunc* | Alignment with identification of exon-exon junctions |
| *propmapped* | Compute mapping statistics |
| *featureCounts* | Compute count matrix for specified genomic features |

Other Rsubread functions are briefly discussed in the Discussion section.

### 3.2 Building the index

*buildIndex* creates a hash table of the target genome from a FASTA file. The index can be built at either single-base or 3-base resolution. Building the full index at single-base resolution takes slightly longer than the gapped index (about 90 minutes vs 18 minutes for the human or mouse genomes) and produces a larger file (about 15Gb vs 5Gb), but allows subsequent alignment to proceed more quickly. The index needs to be built only once for each genome so single-based resolution is set as the default choice in Rsubread versions 1.30.4 and later. Building a gapped index may however be an efficient choice for smaller projects and was the default in older versions of the software.

### 3.3 Alignment

Alignment itself is performed by either the *align* or *subjunc* functions. Both functions accept raw reads, in the form of Fastq, SAM or BAM files, and output read alignments in either SAM or BAM format.

The *align* function is exceptionally flexible. It performs local read alignment and reports the largest mappable region for each read, with unmapped read bases being soft-clipped. Its unique seed-and-vote design makes it suitable for RNA-seq as well as for genomic DNA sequencing experiments. It automatically detects insertions and deletions. *align* is recommended for gene-level expression analyses of RNA-seq or for any type of DNA sequencing.

The *subjunc* function is similar to *align* but provides comprehensive detection of exon-exon junctions and reports full alignments of intron-spanning reads. *subjunc* is recommended for any RNA-seq analysis requiring intra-gene resolution.

Both *align* and *subjunc* achieve high accuracy via a two-pass process. The first pass is the seed-and-vote step, by which a large number of 16mer subreads from each read are mapped to the genome using the hash table. This step detects indels and exon-exon junctions and determines the major mapping location of the read. The second pass undertakes a detailed local re-alignment of each read with the aid of collected indels and junctions. Subread and Subjunc were the first aligners to implement such a two-pass strategy [13].

*align* and *subjunc* support reads from any of the major Next Generation Sequencing (NGS) technologies. Users can specify the amount of computer memory and the number of threads to be used, enabling the aligners to run efficiently on a variety of computer hardwares from super-computers to personal computers.

The *propmapped* function calculates the proportion of reads or fragments that are successfully mapped, a useful quality assessment metric.

### 3.4 Counting reads

The *featureCounts* function counts the number of reads or read-pairs that overlap any specified set of genomic features. It can assign reads to any type of genomic region. Regions may be specified as simple genomic intervals, such as promoter regions, or can be collections of genomic intervals, such as genes comprising multiple exons. Any set of genomic features can be specified in GTF, GFF or SAF format, either as a file or as an R data.frame. SAF is a Simplified Annoation Format with columns GeneID, Chr, Start, End and Strand.

*featureCounts* produces a matrix of genewise counts suitable for input to gene expression analysis packages such as limma [7], edgeR [9] or DESeq2 [10]. Alternatively, a matrix of exon-level counts can be produced suitable for differential exon usage analyses using limma, edgeR or DEXSeq [4].

*featureCounts* outputs the genomic length and position of each feature as well as the read count, making it straightforward to calculate summary measures such as RPKM (reads per kilobase per million reads).

*featureCounts* includes a large number of powerful options that allow it to be optimized for different applications. Reads that overlap more than one feature can be ignored, multi-counted or counted fractionally. Reads can be extended before counting to allow for probable fragment length. Minimum overlap or minimum quality score metrics can be specified. Reads can be counted in a strand specific or non-specific manner. Reads that span exon-exon junctions can be counted, as can reads that are internal to exons.

### 3.5 Counting reads at the exon level

RNA-seq data can be used not only for gene expression but also to investigate alternative use of exons occurring during transcription of genes. To detect alternative exon use, the abundance of exons needs to be accurately measured. Of particular importance is the mapping and counting of exon-spanning reads that span two or more exons in the same gene. Exon-spanning reads typically account for around 20 to 30 percent of reads in an RNA-seq dataset.

To use Rsubread for exon-level RNA-seq analysis, the *subjunc* aligner should be used for read mapping as it performs full alignment for each read. Reads mapped by *subjunc* can then be assigned to exons using *featureCounts*, which should be run at the feature level to allow reads to be assigned to exons instead of genes. Multi-overlapping reads should be counted so that exon-spanning reads can be assigned to all their overlapping exons.

### 3.6 Built-in annotation

Rsubread comes with annotation for human and mouse genes already installed, so that GTF or SAF files do not need to be specified for these species. The built-in annotation follows NCBI RefSeq gene annotation with the simplification that overlapping exons from the same gene are merged. This simplification reduces annotation ambiguity and proves beneficial for most RNA-seq expression analyses. Built-in annotation is provided for the mm9, mm10, hg19 and hg38 genome builds. Rsubread’s built-in hg38 annotation was used for the simulations and comparisons reported in this article.

### 3.7 Quantification at the gene-level: speed and memory

We now compare the Rsubread gene-level workflow, which consists of *align* and *featureCounts*, to other workflows that generate read counts for genes. Rsubread is compared to aligners TopHat2 [25] and STAR [26] combined with quantification tools htseq-count [3], *summarizeOverlaps* and *featureCounts*. Google Scholar searches suggest that these are currently the most popular tools for generating gene-level counts. htseq-count is part of the HTSeq Python library. *summarizeOverlaps* is a function in the Bioconductor package GenomicRanges.

First we assessed the running time of the read aligners on the SEQC UHRR sample (Figure 1). *align* was slightly faster when run with the full genome index (*align*-F) as opposed to the gapped index (*align*-G), with both options being faster than STAR or TopHat2. STAR was 50% slower than *align*-F and TopHat2 more than an order of magnitude slower.

**Figure 1:**
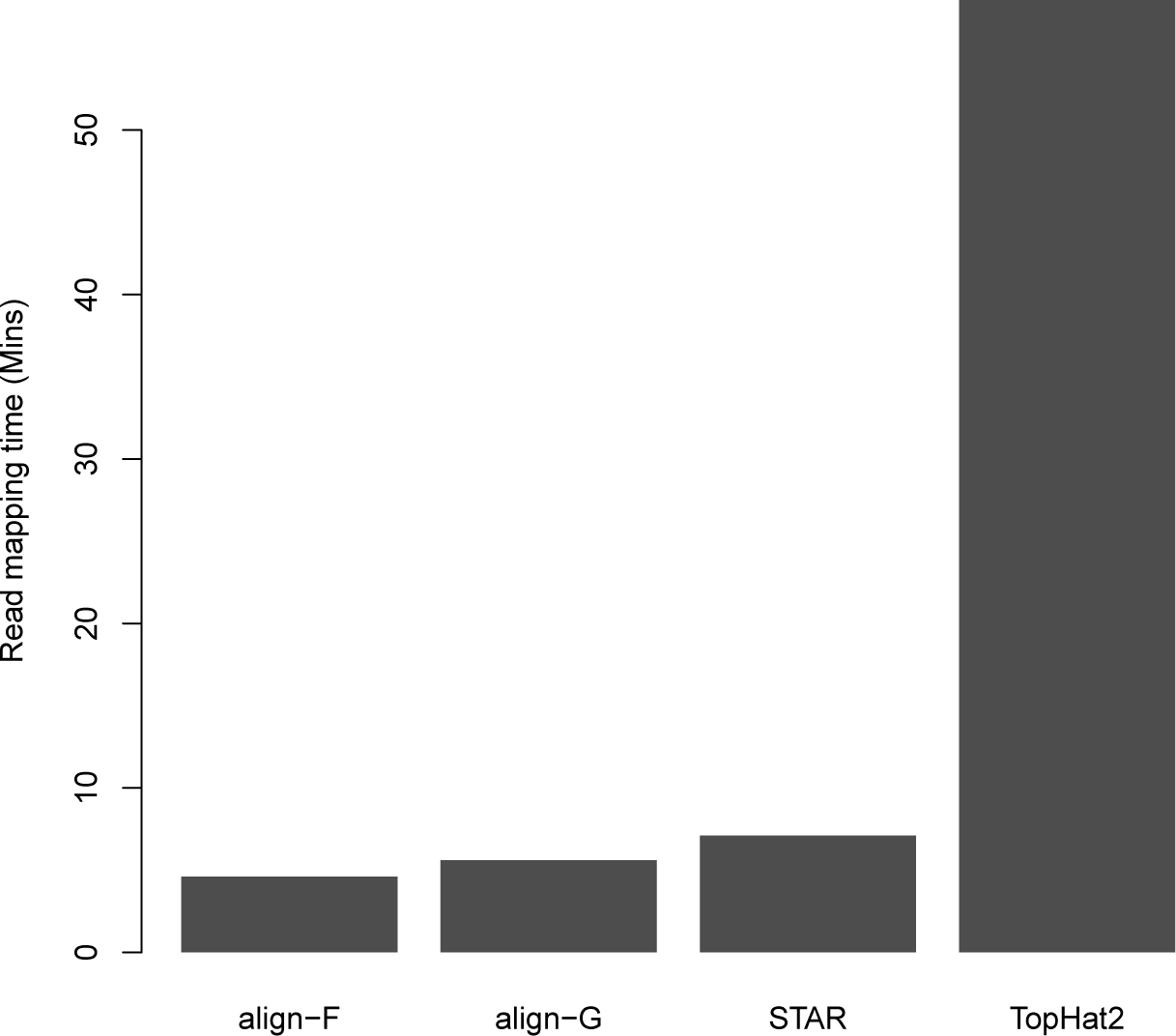
Run times of read aligners. Each aligner used ten threads to map 15 million 100bp read-pairs from the SEQC UHRR sample to the human reference genome GRCh38. Rsubread::align is faster than STAR or TopHat2 regardless of whether the full index (align-F) or a gapped index (align-G) is used.

TopHat2 and *align*-G had the smallest memory footprints for the same operation (supplementary Figure S1). *align*-F used twice as much memory and STAR over four times as much.

Next we ran the quantification tools to assign the mapped UHRR reads to RefSeq human genes. This showed *featureCounts* to be 16 – 175 times faster than the other tools (Figure 2). *featureCounts* was equally as fast regardless of the alignment used. *summarizeOverlaps* and htseq-count were slower when working on the TopHat2 alignment than the STAR alignment. In this evaluation, *align* and STAR output name-sorted aligned reads whereas TopHat2 output location-sorted reads.

**Figure 2:**
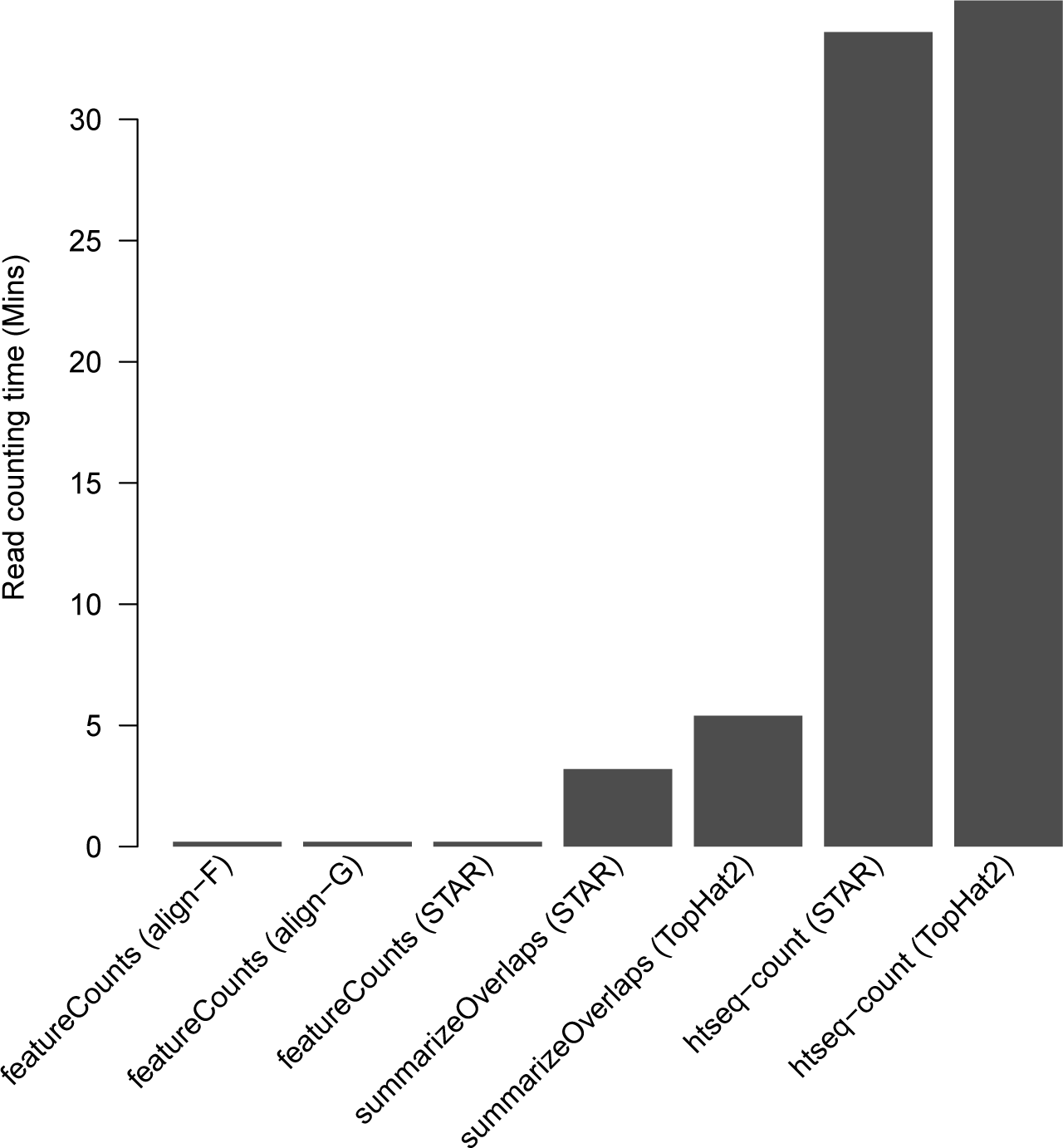
Running time of different quantification tools. Labels under each bar indicate the quantification method and the aligner (in parenthesis) that produced the mapped reads used for counting. Mapped reads were assigned to NCBI RefSeq human genes. *FeatureCounts* is the only tool that supports multi-threaded read counting and it was run with four threads.

*featureCounts* used easily the least memory for the quantification step (supplementary Figure S2). htseq-count used only slightly more memory than *featureCounts* when read pairs were name-sorted, but it used *>*40 times more memory when the read pairs were location-sorted. *summarizeOverlaps* had high memory usage for both name-sorted and location-sorted reads because it loads all the reads into memory at once.

In summary, Rsubread outperformed the STAR-based workflows in both speed and memory use. While TopHat2 has a slightly smaller memory footprint than Rsubread for alignment, it was far too slow to be competitive.

### 3.8 Quantification at the gene-level: accuracy

Next we compared workflows for accuracy in quantifying gene expression levels. First we ran the workflows on the UHRR and HBRR samples from the SEQC Project. Read counts generated from each workflow were then compared to the expression levels of 958 genes as measured by TaqMan RT-PCR, a high-throughput quantitative PCR technique. RNA-seq counts were converted to log2-FPKM (fragments per kilobases per million) values and TaqMan RT-PCR data were also converted to log2 scale.

Rsubread workflows are found to yield the highest correlation with TaqMan RT-PCR data for both UHRR and HBRR samples (Table 2). All workflows produce higher correlation for HBRR sample than for UHRR sample, which is expected because the UHRR sample is made up from multiple cancer cell lines.

**Table 2:** Gene-level accuracy comparison. The table gives Pearson correlations between true log2-expression levels and log2-FPKM values produced by each workflow. The align-F + featureCounts workflow gives the best correlation in each case.

| Workflow | UHRR | HBRR | Simulation |
| --- | --- | --- | --- |
| align-F + featureCounts | **0.851** | **0.870** | **0.955** |
| align-G + featureCounts | 0.850 | 0.869 | **0.955** |
| STAR + featureCounts | 0.848 | 0.867 | 0.901 |
| STAR + htseq-count | 0.845 | 0.864 | 0.877 |
| STAR + summarizeOverlaps | 0.845 | 0.864 | 0.877 |
| TopHat2 + htseq-count | 0.843 | 0.863 | 0.921 |
| TopHat2 + summarizeOverlaps | 0.843 | 0.863 | 0.921 |

Columns “UHRR” and “HBRR” are for the SEQC UHRR and SEQC HBRR samples respectively. For the SEQC columns, the log2-expression values of 958 genes measured by TaqMan RT-PCR are taken as true values. Column “Simulation” shows simulation results for 28,395 genes. For all columns, an offset of 0.25 was added to raw gene counts to avoid taking logarithms of zeros.

Next we compared the workflows on simulated data. All the workflows were run on a simulated FASTQ file of 15 million read pairs. Read counts were converted to log2-FPKM and compared to the known log2-FPKM values from which the simulated sequence reads were generated. The Rsubread workflows are again found to achieve the best correlation with the true expression values (Table 2). Both Rsubread workflows yield correlation > 0.95, much higher than those for other workflows.

In general, *align* was more accurate than STAR and *featureCounts* was more accurate than either htseq-count or *summarizeOverlaps. summarizeOverlaps* uses the same counting strategy as that developed by htseq-count and therefore gives the same results.

### 3.9 Quantification at the exon level: speed and memory

Next we compared workflows to obtain exon-level read counts. Rsubread workflows for exon-level analysis comprise the *subjunc* program, which was run with a full genome index (*subjunc*-F) or a gapped index (*subjunc*-G), and the *featureCounts* programs. Rsubread was compared to TopHat2 + dexseq count, TopHat2 + *countOverlaps*, STAR + dexseq count, STAR + *countOverlaps* and STAR + *featureCounts*. dexseq count.py is a Python script that comes with the DEXSeq package for counting RNA-seq reads by exon. *countOverlaps* is a function in IRanges package. For all pipelines, reads spanning multiple exons were counted for all the relevant exons.

All workflows were run on the SEQC UHRR data. For read mapping, *subjunc*-F was the fastest, followed by STAR, *subjunc*-G and TopHat2 (supplementary Figure S3). STAR and *subjunc*-G were almost equal while *subjunc*-F was about 20% faster. TopHat2 was about 10 times slower.

For read counting, *featureCounts* is more than an order of magnitude faster than countOverlaps or dexseq count, regardless of which aligner output was used (supplementary Figure S4). Dexseq count was the slowest counting tool.

*subjunc* uses much less memory than STAR (supplementary Figure S5) and *featureCounts* uses less memory than dexseq count or countOverlaps (supplementary Figure S6). As for the gene-level results, TopHat2 used slightly less memory than *subjunc*-G but at too high a price in terms of running time.

In summary, *subjunc*-F and *featureCounts* constitute the fastest workflow for exon-level analysis of RNA-seq data. *featureCounts* uses the least memory of any quantification tool and *subjunc* uses less memory than STAR.

### 3.10 Quantification at the exon level: accuracy

We used the same simulated data as for the gene-level comparison to assess the accuracy of the exon-level workflows. Overlapping exons found between genes were removed from analysis to avoid counting ambiguity. Exons from genes appearing in more than one chromosome, or appearing in both strands of the same chromosome, were also removed because dexseq count cannot process such exons. 5000 exons were excluded from this analysis in total (out of 261,752 exons). Read counts from remaining exons were then converted to log2-FRKMs for comparison.

As was seen in the gene-level comparison, the two Rsubread workflows both out-performed the other workflows (Table 3). *subjunc* and TopHat2 were more accurate than STAR. *featureCounts* was more accurate than *countOverlaps* and *countOverlaps* was more accurate than dexseq count. The accuracy of the workflows was affected by both read mapping and counting. The STAR + dexseq count workflow had the worst correlation of the workflows. Replacing dexseq count with *featureCounts* improved the accuracy, but it remained lower than that for the two pure Rsubread workflows.

**Table 3:** Exon-level accuracy comparison. The table shows the Pearson correlation between the true log2-FPKM expression values of exons and log2-FPKM values produced by each workflow. Rsubread workflows give the best correlation with true values.

| Workflow | Correlation |
| --- | --- |
| subjunc-F + featureCounts | **0.982** |
| subjunc-G + featureCounts | **0.982** |
| STAR + featureCounts | 0.950 |
| STAR + dexseq_count | 0.927 |
| STAR + countOverlaps | 0.950 |
| TopHat2 + dexseq_count | 0.942 |
| TopHat2 + countOverlaps | 0.980 |

A offset of 0.25 was added to raw exon counts to avoid taking logarithms of zero values.

### 3.11 Detection and quantification of exon-exon junctions

Exon-exon junctions can be discovered directly by the correct mapping of junction-spanning reads (junction reads). Most RNA-seq aligners report the locations of exon splice sites (donor and receptor sites). The number of reads supporting the splice sites detected are often reported as well, providing a quantitive measurement for the junctions events. Analysis of discovered junctions and their abundance is an important step in the discovery of alternative splicing events. Junction data can be further combined with exon-level and gene-level expression data in order to detect differentially spliced genes.

Nevertheless, junction reads can be difficult to map correctly because they may span an intron tens of thousands of bases long and indeed might span more than one intron. Here we assess the performance of the aligners for mapping junction reads and calling junctions. We used the same simulated data as above. There are 233,021 exon-exon junctions in the simulated data and 25% of the simulation reads are junction reads.

*subjunc*-F and *subjunc*-G had better overall performance than STAR or TopHat2 as measured by the F1 summary of precision and recall (Table 4). In particular, *subjunc*-F and *subjunc*-G outperformed STAR and TopHat2 in mapping of junction reads by a clear margin. STAR was slightly less sensitive than the other aligners in mapping junctions reads or detecting junctions.

**Table 4:** Aligner performance in mapping junction reads and reporting exon-exon junctions. Results are based on simulated data.

| Workflow | Junctions |  |  | Junction reads |  |  |
| --- | --- | --- | --- | --- | --- | --- |
|  | Recall | Prec | F1 | Recall | Prec | F1 |
| subjunc-F | 99.80 | 99.53 | **99.66** | 95.51 | 98.18 | **96.82** |
| subjunc-G | 99.82 | 99.43 | 99.63 | 95.50 | 98.16 | 96.81 |
| STAR | 98.48 | 99.87 | 99.17 | 89.87 | 98.06 | 93.78 |
| TopHat2 | 99.12 | 99.91 | 99.51 | 90.15 | 98.57 | 94.17 |

Column “Recall” gives the percentage of correctly called junctions (or junction reads) out of all junctions (or junction reads) generated in the simulated dataset. Column “Precision” gives the percentage of correctly called junctions (or junction reads) out of all reported junctions (or junction reads). Column “F1” gives the F1 score that is the harmonic mean of precision and recall.

## 4 Discussion

Read mapping and quantification are computationally-intensive operations that lay the foundation for most analyses of NGS data. The time-consuming and resource-hungry nature of these operations is a major bottleneck for larger projects. Meanwhile, R is an easy-to-learn scripting language that is widely used for statistical analyses of NGS data once the processing of the raw reads is completed. Rsubread provides functions for read mapping and quantification within the R programming environment, allowing an entire NGS analysis, from reads to results, to be completed in a single R session. Rsubread can work with Bioconductor packages limma, edgeR and DESeq2 to complete an entire RNA-seq analysis in R from read mapping through to the discovery of genes that exhibit significant expression changes [21, 22]. It has proved a valuable resource for NGS analyses in biomedical research [27].

As well as ease-of-use, this study has shown that Rsubread outperforms the most popular competing alignment and quantification tools regardless of programming language. Rsubread was generally found to be faster, to use less memory and to provide more accurate expression quantification than the competitor tools. The improved accuracy of the Rsubread workflows should translate into more accurate downstream analyses such as discovery of differentially expressed genes. The main exception is that TopHat2 used slightly less memory for alignment than Rsubread. TopHat2 however was an order of magnitude slower and hence was not considered competitive overall. Our comparisons included only the two or three most popular RNA-seq aligners and the two or three most popular counting tools and are not intended to be a comprehensive study of all current RNA-seq tools. Nevertheless, the comparisons are representative of current practice and are sufficient to show that users can adopt the R environment for alignment and quantification without suffering any loss of performance.

The Rsubread package continuously tracks the stand-alone C programs provided by the Subread project (http://subread.sourceforge.net), meaning that the R functions always give equivalent performance to the corresponding C programs. Users of R can therefore take advantage of the Rsubread user-interface without compromising performance.

We chose STAR and TopHat2 for the comparisons in this article because they remain easily the most popular RNA-seq aligners in the published literature. The TopHat website has recently recommended, however, that users migrate to HISAT2 instead. Our comparisons show that HISAT2 is indeed fast and economical with memory but is less able than the other aligners to detect exon-exon junctions (data not shown). Rsubread outscores HISAT2 on all the accuracy criteria presented in this article (data not shown).

Although the main focus of this study is on the analysis of RNA-seq data, Rsubread can be used also for the analysis of other types of sequencing data such as histone ChIP-seq and ATAC-seq. The *align* function can be used for read mapping and *featureCounts* can be used to produce read counts for promoter regions or gene bodies to provide a measurement of peak abundance [28, 29].

The Rsubread package includes other functions beyond the scope of this article including *sublong* (for long read mapping), *exactSNP* (SNP identification), *atgcContent* (compute nucleotide frequencies), *detectionCall*, and *promoterRegions*.

## 6 Acknowledgements

The authors thank Jenny Dai and Timothy Triche, Jr., for code contributions to Rsubread.

## References

[1] Liao, Y., Smyth, G.K. and Shi, W. (2014) featureCounts: an efficient general-purpose read summarization program. Bioinformatics, 30, 923–930.

[2] Su, Z., Labaj, P.P., Li, S., Thierry-Mieg, J., Thierry-Mieg, D., Shi, W., Wang, C., Schroth, G.P., Setterquist, R.A., Thompson, J.F. et al. (2014) A comprehensive assessment of RNA-seq accuracy, reproducibility and information content by the Sequencing Quality Control Consortium. Nature Biotechnology, 32, 903–914.

[3] Anders, S., Pyl, P.T. and Huber, W. (2015) HTSeq—a Python framework to work with high-throughput sequencing data. Bioinformatics, 31, 166–169.

[4] Anders, S., Reyes, A. and Huber, W. (2012) Detecting differential usage of exons from RNA-seq data. Genome Research, 22, 2008–2017.

[5] R Core Team (2018) R: A Language and Environment for Statistical Computing. R Foundation for Statistical Computing, Vienna, Austria.

[6] Huber, W., Carey, V.J., Gentleman, R., Anders, S., Carlson, M., Carvalho, B.S., Bravo, H.C., Davis, S., Gatto, L., Girke, T. et al. (2015) Orchestrating high-throughput genomic analysis with Bioconductor. Nature Methods, 12, 115–121.

[7] Ritchie, M.E., Phipson, B., Wu, D., Hu, Y., Law, C.W., Shi, W. and Smyth, G.K. (2015) limma powers differential expression analyses for RNA-sequencing and microarray studies. Nucleic Acids Research, 43, e47.

[8] Law, C.W., Chen, Y., Shi, W. and Smyth, G.K. (2014) Voom: precision weights unlock linear model analysis tools for RNA-seq read counts. Genome Biology, 15, R29.

[9] Robinson, M., McCarthy, D. and Smyth, G. (2010) edgeR: a Bioconductor package for differential expression analysis of digital gene expression data. Bioinformatics, 26, 139–140.

[10] Love, M.I., Huber, W. and Anders, S. (2014) Moderated estimation of fold change and dispersion for RNA-seq data with DESeq2. Genome Biol., 15, 550.

[11] Trapnell, C., Pachter, L. and Salzberg, S.L. (2009) TopHat: discovering splice junctions with RNA-seq. Bioinformatics, 25, 1105–1111.

[12] Dobin, A., Davis, C.A., Schlesinger, F., Drenkow, J., Zaleski, C., Jha, S., Batut, P., Chaisson, M. and Gingeras, T.R. (2013) STAR: ultrafast universal RNA-seq aligner. Bioinformatics, 29, 15–21.

[13] Liao, Y., Smyth, G.K. and Shi, W. (2013) The Subread aligner: fast, accurate and scalable read mapping by seed-and-vote. Nucleic Acids Research, 41, e108.

[14] Kim, D., Langmead, B. and Salzberg, S.L. (2015) HISAT: a fast spliced aligner with low memory requirements. Nature Methods, 12, 357.

[15] Quinlan, A. and Hall, I. (2010) BEDTools: a flexible suite of utilities for comparing genomic features. Bioinformatics, 26, 841–842.

[16] Schmid, M.W. and Grossniklaus, U. (2015) Rcount: simple and flexible RNA-Seq read counting. Bioinformatics, 31, 436–437.

[17] Gaidatzis, D., Lerch, A., Hahne, F. and Stadler, M.B. (2015) QuasR: quantification and annotation of short reads in R. Bioinformatics, 31, 1130–1132.

[18] Langmead, B., Trapnell, C., Pop, M. and Salzberg, S.L. (2009) Ultrafast and memory-efficient alignment of short DNA sequences to the human genome. Genome Biology, 10, Article R25.

[19] Au, K.F., Jiang, H., Lin, L., Xing, Y. and Wong, W.H. (2010) Detection of splice junctions from paired-end RNA-seq data by SpliceMap. Nucleic Acids Research, 38, 4570–4578.

[20] Döring, A., Weese, D., Rausch, T. and Reinert, K. (2008) SeqAn an efficient, generic C++ library for sequence analysis. BMC Bioinformatics, 9, 11.

[21] Chen, Y., Lun, A.T.L. and Smyth, G.K. (2016) From reads to genes to pathways: differential expression analysis of RNA-seq experiments using Rsubread and the edgeR quasi-likelihood pipeline. F1000Research, 5, 1438.

[22] Schmid, M.W. (2017) RNA-Seq data analysis protocol: Combining in-house and publicly available data. Methods in Molecular Biology, 1669, 309–335.

[23] Engstrom, P.G., Steijger, T., Sipos, B., Grant, G.R., Kahles, A., Ratsch, G., Goldman, N., Hubbard, T.J., Harrow, J., Guigo, R. et al. (2013) Systematic evaluation of spliced alignment programs for RNA-seq data. Nature Methods, 10, 1185–1191.

[24] Baruzzo, G., Hayer, K.E., Kim, E.J., Camillo1, B.D., FitzGerald, G.A. and Grant, G.R. (2017) Simulation-based comprehensive benchmarking of RNA-seq aligners. Nature Methods, 14, 135–139.

[25] Kim, D., Pertea, G., Trapnell, C., Pimentel, H., Kelley, R. and Salzberg, S.L. (2013) TopHat2: accurate alignment of transcriptomes in the presence of insertions, deletions and gene fusions. Genome Biol, 14, R36.

[26] Dobin, A., Davis, C.A., Schlesinger, F., Drenkow, J., Zaleski, C., Jha, S., Batut, P., Chaisson, M. and Gingeras, T.R. (2013) STAR: ultrafast universal RNA-seq aligner. Bioinformatics, 29, 15–21.

[27] Fong, C.Y., Gilan, O., Lam, E.Y., Rubin, A.F., Ftouni, S., Tyler, D., Stanley, K., Sinha, D., Yeh, P., Morison, J. et al. (2015) BET inhibitor resistance emerges from leukaemia stem cells. Nature, 525, 538–542.

[28] Pal, B., Bouras, T., Shi, W., Vaillant, F., Sheridan, J., Fu, N., Breslin, K., Jiang, K., Ritchie, M., Young, M. et al. (2013) Global changes in the mammary epigenome are induced by hormonal cues and coordinated by Ezh2. Cell Reports, 3, 411–426.

[29] de Santiago, I. and Carroll, T. (2018) Analysis of ChIP-seq data in R/Bioconductor. Methods in Molecular Biology, 1689, 195–226.

